# The Rayleigh Quotient and Contrastive Principal Component Analysis II

**DOI:** 10.64898/2026.04.08.717236

**Authors:** Kayla Jackson, Maria Carilli, Lior Pachter

**Affiliations:** Division of Biology and Biological Engineering, California Institute of Technology, Pasadena, CA, USA; Department of Computing and Mathematical Sciences, California Institute of Technology, Pasadena, CA, USA; Keck School of Medicine, University of Southern California, Los Angeles, CA, USA

**Author notes:** Authors contributed equally.

## Abstract

Contrastive principal component analysis (PCA) methods are effective approaches to dimensionality reduction where variance of a target dataset is maximized while variance of a background dataset is minimized. We previously described how contrastive PCA problems can be written as solutions to generalized eigenvalue problems that maximize particular instantiations of the Rayleigh quotient. Here, we discuss two extensions of contrastive PCA: we use kernel weighting from spatial PCA (k-*ρ*PCA) to contrast spatial and non-spatial axes of variation, and separately solve the Rayleigh quotient in the space of basis function coefficients (f-*ρ*PCA) to find modes of variation in functional data. Together, these extensions expand the scope of contrastive PCA while unifying disparate fields of spatial and functional methods within a single conceptual and mathematical framework. We showcase the utility of these extensions with several examples drawn from genomics, analyzing gene expression in cancer and immune response to vaccination.

## Introduction

Contrastive PCA methods are an approach to dimensionality reduction that simultaneously maximizes variance in a target dataset while minimizing the variance in a background dataset. Our recent work has shown that this objective can be formulated as a Rayleigh quotient maximization problem and solved by means of a generalized eigenvalue problem (Carilli et al., 2025). Here, we show that the method, called *ρ*PCA, can be extended to yield contrastive analogues of spatial and functional PCA.

Kernel PCA (kPCA) is an extension of PCA that enables identification of complex and nonlinear patterns in multi-dimensional data (Schölkopf et al., 1997). Kernel-based methods rely on mapping the input data to a (possibly infinite-dimensional) Hilbert space 𝒱 using a feature map *ϕ* : 𝒳 → 𝒱. In this framework, kPCA performs eigendecomposition of the covariance operator defined as

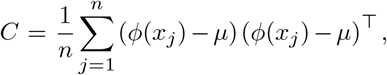

where 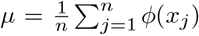. Importantly, practical applications of kPCA do not require an explicit form of *ϕ*, as any positive semi-definite kernel corresponds to a map into a Hilbert space 𝒱 such that the kernel function computes the inner product in that space.

Recent methods for performing spatial PCA have proposed the use of spatial kernels in a latent factor model or the eigendecomposition of adjacency-weighted sample covariance matrices to reflect the underlying structure across the spatial domain (Shang and Zhou, 2022; Jombart et al., 2008). We propose a contrastive spatial PCA by replacing the standard covariance matrix with its kernel-weighted counterpart in *ρ*PCA to facilitate spatially-weighted Rayleigh quotient problems. This extension leverages kernels to capture localized spatial structure while maintaining the contrastive advantages of *ρ*PCA.

Functional PCA (fPCA) is a technique that extends PCA to reduce the dimensionality of data that can be represented as curves or functions over a continuous domain (e.g., time or space) by finding dominant “modes of variation” (Ramsay and Silverman, 2002, 2005). In this setting, *n* independent observations *X*_*i*_(*t*), …, *X*_*n*_(*t*) are considered realizations of a square-integrable stochastic process *X*(*t*) over the continuous variable *t*. Let *µ*(*t*) = *E*[*X*(*t*)] and *c*(*s, t*) = Cov(*X*(*s*), *X*(*t*)) = *E*[(*X*(*s*) −*µ*(*s*))(*X*(*t*) −*µ*(*t*))] be the mean function and covariance function of the process *X*(*t*), respectively. The covariance operator *C* : *L*^2^(𝒯) → *L*^2^(𝒯) acts on square-integrable functions *f* (*s*) ∈ *L*^2^(𝒯) as

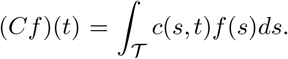

As *C* is compact, self-adjoint (symmetric, or *c*(*s, t*) = *c*(*t, s*)), and positive semi-definite (∫_𝒯_ ∫_𝒯_ *f*(*s*)*c*(*s, t*)*f* (*t*)*dsdt*≥ 0), by Mercer’s theorem, the function *c*(*s, t*) can be expressed as

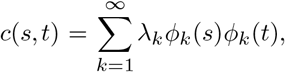

where the *ϕ*_*k*_ are the eigenfunctions of *C* satisfying *Cϕ*_*k*_ = *λ*_*k*_*ϕ*_*k*_ and *λ*_1_ ≥ *λ*_2_ ≥ … ≥ 0. The orthonormality of eigenfunctions requires ∫_𝒯_ *ϕ*_*k*_(*t*)*ϕ*_*l*_(*t*)*dt* = 0 for *k* ≠ *l* and ∫_𝒯_ *ϕ*_*k*_(*t*)*ϕ*_*l*_(*t*)*dt* = 1 for *k* = *l*.

fPCA seeks to find orthogonal functions that capture the most variation in *X*(*t*), or, analogously to PCA, maximize the variance of the integral inner product (the process projected onto the new function):

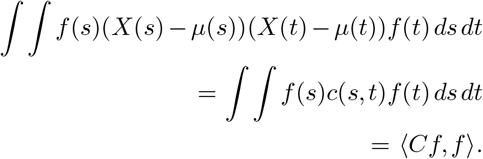

Recalling the eigendecomposition of *C*, the functions that capture the most variation are the eigenfunctions *ϕ*_*k*_(*t*), which form a basis for *L*^2^ such that each process *X*_*i*_(*t*) can be written as

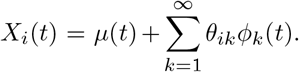

Each observation *i* has weights *θ*_*ik*_ associated with each eigenfunction *k* that capture how strongly the eigenfunction contributes to the observation.

Practically, fPCA is implemented in one of two ways. In the first approach, observations are sampled on a grid of points over the continuous variable, and traditional PCA is employed on the resulting matrices. The eigenvectors can be interpolated between discrete features to produce eigenfunctions over a continuous feature space (Ramsay and Silverman, 2005). In the second approach, samples are first represented as combinations of a finite set of basis functions (e.g., B-splines, monomials, or Fourier coefficients). PCA is then performed on the coefficients of the bases (taking into account the non-orthogonality of basis functions if necessary) to find eigenvector coefficients for the set of basis functions (Ramsay and Silverman, 2005; Wang et al., 2016).

fPCA has been adapted to various settings. For example, it has been used to fit observations sampled at irregular or sparse time-points by smoothing covariance and mean functions (Yao et al., 2005), combined with mixed-effect models (James et al., 2000), and applied to adjust for known covariates (Jiang and Wang, 2010). A recent attempt to implement a contrastive version with target and background functional data has been proposed that mimics the contrastive PCA method of (Abid et al., 2018) by subtracting from the target covariance function a weighted version of the background covariance function (Zhang and Li, 2025). This approach to contrastive fPCA inherits the same problems as the (Abid et al., 2018) contrastive PCA method (Carilli et al., 2025), including the issue that a difference of positive semi-definite matrices may not be positive semi-definite, and requiring a tunable contrastive parameter that can produce arbitrary eigenfunctions (see Supp. Fig. S1). We show that contrastive functional dimension reduction can be achieved more naturally by performing *ρ*PCA in the space of basis coefficients, as shown in Figure 2A, leading to meaningful modes of variation present in target curves and absent in background curves.

## Results

### k-*ρ*PCA Application to Visium and VisiumHD

We first applied k-*ρ*PCA (Fig. 1A) to a dataset containing colorectal cancer (CRC) and nearby non-tumor tissue samples profiled with Xenium, Visium V2, Visium HD, and Chromium Single Cell Gene Expression Flex (Oliveira et al., 2025). We focused on data from Patient 2 that had Visium V2 and Visium HD CRC samples available for use as “target” (Fig. 1B) and a non-spatial single cell RNA-seq (scRNA-seq) data from non-tumor adjacent tissue available to use as “background.” The k-*ρ*PCA method directly identifies spatially variable genes in the cancer sample while accounting for normal cell type variation.

**Fig. 1.**
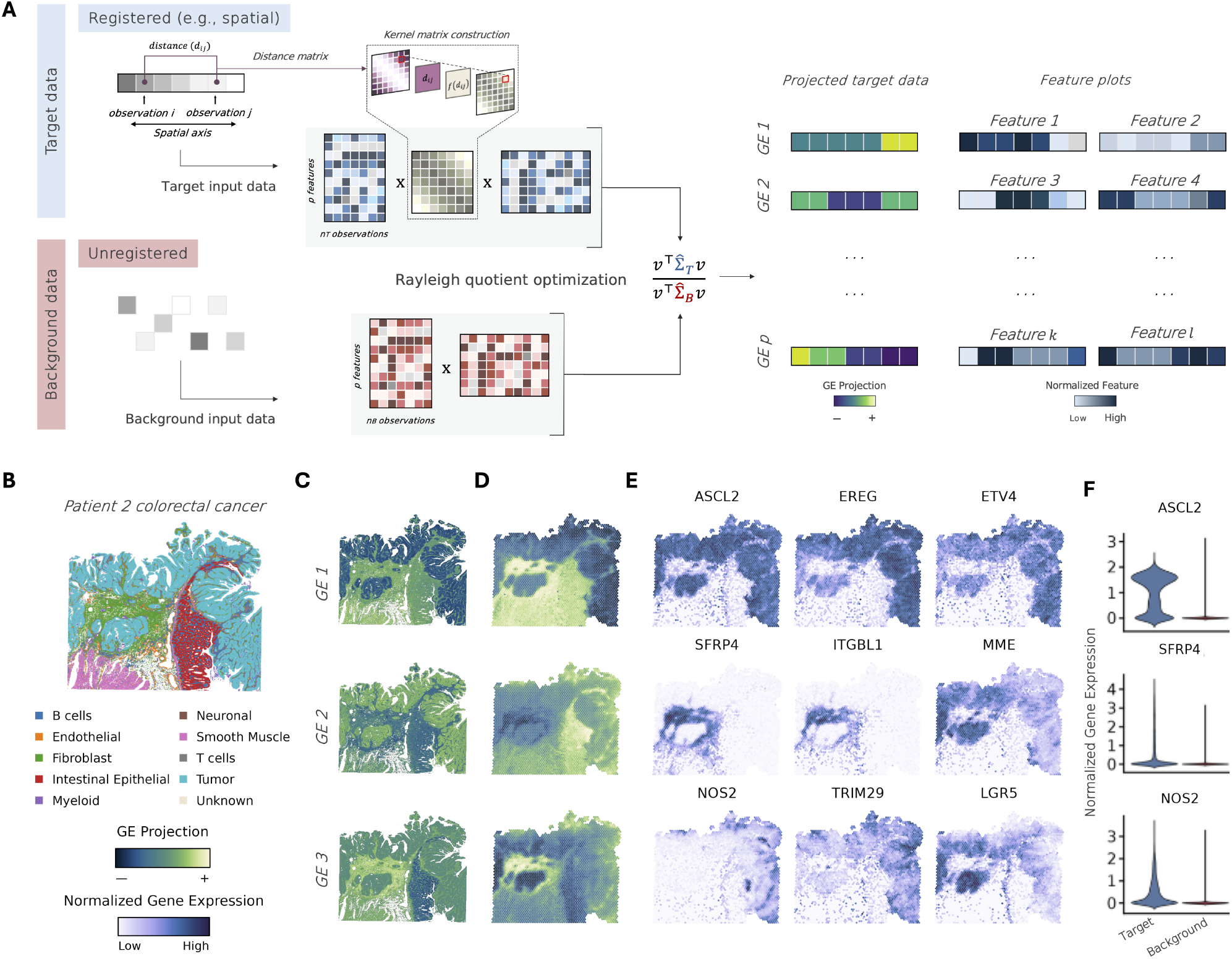
**A**. As in *ρ*PCA, k-*ρ*PCA is useful in settings where the goal is to maximize variance in a “target” dataset of interest relative to a “background” dataset. However, in k-*ρ*PCA, the target dataset is spatially or temporally registered, which enables the computation of a kernel matrix that weights pairs of observations using a function of the spatial (temporal) distance between them. The kernel matrix is subsequently used to weight observations in the target covariance matrix, 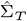. In contrast, the background covariance matrix, 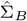, is estimated from unregistered data. The generalized eigenvectors that maximize the Rayleigh quotient define the directions that maximize variance in the target and minimize variance in the background. The target data can then be projected onto these directions and visualized in space (time), together with the features that exhibit large-magnitude loadings on each generalized eigenvector. In this example, we applied k-*ρ*PCA to a spatial colorectal tumor sample (**B**.) and used non-spatial data from nearby non-tumor tissue as the background. The tumor sample was profiled with (**C**.) Visium HD and (**D**.) Visium V2. In **E**., genes with high magnitude along the first few GEs computed from the Visium V2 data highlight tumor-normal boundaries and specific tumor compartments. **F**. Selected gene expression plots showing greater overall variance in the target dataset.

Our results show that generalized eigenvector (GE) 1 clearly differentiates tumor and non-tumor-labeled spots in both Visium HD (Fig. 1C) and Visium (Fig. 1D). This is in contrast to standard PCA which suggests that the non-tumor intestinal epithelium and tumor are similar in the dominant direction of variation (Supp. Fig. S2). Importantly, GE 1 distinguishes tumor and non-tumor tissue even when an un-matched sample is used as a background (Supp. Fig. S3).

Exploration of the genes associated with each GE can provide insight into the progression and development of CRC (Fig. 1E). For instance, genes such as *ASCL2, EREG*, and *SFRP* have been previously associated with tumor prognosis, metastasis, and disease progression in CRC and other cancers (Zhang et al., 2023; Fonseca et al., 2021). Although this analysis specifically identifies genes with high spatial variance, we also find that, in general, these genes are more variable in cancer than in normal adjacent tissue (Fig. 1F). Interestingly, two genes associated with GE 2 highlight a fibroblast response in the internal tumor compartment. *ITGBL1* is involved in cell-extracellular matrix adhesion and has been frequently associated with cancer metastasis, while *SFRP4* is a modulator of Wnt signaling and its overexpression in CRC has been associated with disease progression and Wnt activation (Fonseca et al., 2021; Han et al., 2006). The localization of these genes to the internal tumor may suggest how dysregulated cells with metastatic potential avoid immune detection. Finally, *NOS2* expression is induced by oxidative stress and has recently been proposed as an early prognosticator of CRC (Xing et al., 2025; Cianchi et al., 2003). The expression of *NOS2* may be closely associated with *TRIM29*, which has been reported to promote the metabolic shift of tumor cells from oxidative phosphorylation to aerobic glycolysis. Together, these examples demonstrate that k-*ρ*PCA effectively reveals tumor-specific mechanisms without requiring substantial metadata or prior cell-type annotation.

### f-*ρ*PCA Application to Longitudinal Bulk RNA-seq

Measurement of blood transcriptomes is an effective way to profile immunological responses before and after vaccination, with the goal of guiding the design of vaccination protocols for best conferred protection and reduction of adverse events (Arunachalam et al., 2021). To demonstrate the ability of f-*ρ*PCA (Fig. 2A) to extract functional patterns from longitudinal data, we applied it to bulk RNA-seq samples collected over a two-week time course before and after a first and second dose of COVID-19 mRNA vaccines (Rinchai et al., 2022). In the study, blood from 23 subjects was sampled immediately before, for 9 days after and 14 days after vaccination (a total of 11 time points) for both an initial “primer” and a secondary “booster” vaccination (Fig. 2B). The original study performed gene set enrichment analyses on the first and second doses separately. They subsequently compared the number and function of significant gene modules to understand how primer and booster responses differed. We applied f-*ρ*PCA to contrast the two time courses jointly, using the first dose as “background” and the second dose as “target,” and identified key genes that differ between the doses.

**Fig. 2.**
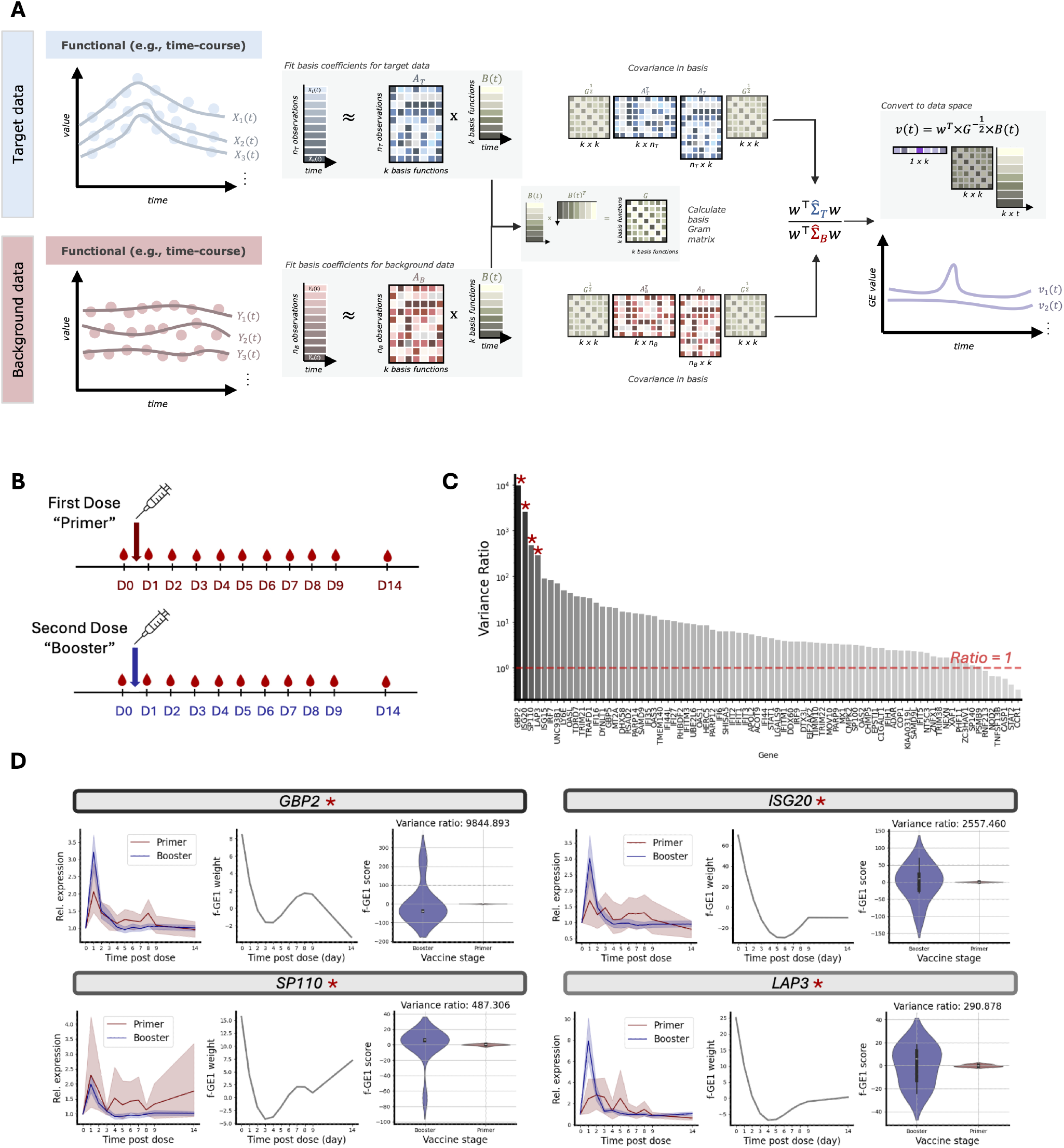
**A**. Diagram of functional *ρ*PCA (f-*ρ*PCA) using a basis representation. Target and background samples are measured at discrete time points, then fit to a set of basis functions to approximate the underlying continuous process. Functional *ρ*PCA calculates the covariance of the fit coefficients, accounting for non-orthogonality of basis functions using the Gram matrix of basis functions, then finds solutions to the Rayleigh quotient. Generalized eigenfunctions are recovered by converting the obtained generalized eigenfunctions back to original data space (e.g., time). See the Methods and Supplementary Methods for details. **B**. We applied f-*ρ*PCA to longitudinal bulk RNA-seq conducted on patients before and after a first (“primer”) and second (“booster”) dose of a COVID-19 mRNA vaccine (diagram adapted from (Rinchai et al., 2022)). **C**. Ratio of the variance of target (“booster”) samples projected onto the first found eigenfunction to background (“primer”) samples projected onto the first eigenfunction for 86 interferon genes. **D**. For four genes with the largest target to background variance ratio, we show the gene expression values (normalized as described in Methods) of primer and booster samples (mean line and shaded standard deviation over the samples indicated), the values over time of the first eigenfunction (f-GE1 weight), and violin plots of the distributions of target and background samples projected onto the first eigenfunction (f-GE1 score). Target samples’ projections display higher variance than background samples.

(Rinchai et al., 2022) report that a set of interferon genes (referred to as module A28 (Altman et al., 2021; Rinchai et al., 2022)) have the most significant response in the first three days after primer administration. Focusing on this set, we obtained fit basis functions to the normalized expression profile for each gene (see Methods). We then performed f-*ρ*PCA on the gene profiles that passed quality control for a total of 86 genes with contrastive analyses (Fig. 2C). The first eigenfunctions have high values at the points in time most variable in the booster that are not variable in the primer (Fig. 2D); for the analyzed interferon genes, there is a peak at the initial administration, consistent with the original paper’s observation that after the second dose “the interferon response was noticeably sharper in comparison to the response observed following the first dose and peaked on day 1 in-stead of day 2” (Rinchai et al., 2022). We also projected fit samples onto the discovered first eigenfunctions for these genes and calculated the ratio of variance of booster samples to primer samples, finding that genes display a range of target and background sample separability (Fig. 2C).

The four genes with the highest booster to primer variance ratio, *GBP2, ISG20, SP110*, and *LAP3*, have previously been associated with SARS-CoV-2 (for expression profiles over all fit samples, see Supp. Fig. S4). *GBP2*, which codes for the GTPase guanylate-binding protein 2, inhibits the cleavage of the SARS-CoV-2 spike protein, thus affecting membrane fusion and viral entry (Mesner et al., 2023). Interferon-stimulated gene 20 (coded for by *ISG20*) is an antiviral RNA exonuclease that has degraded RNA vectors derived from SARS-CoV-2 replicons (Furutani et al., 2021). *SP110* was identified as significantly associated with extreme COVID-19 phenotypes (Mir et al., 2025); and *LAP3*, shown to have increased expression during COVID-19 infection (Witkowski et al., 2021), also exhibits poly(A)-tail elongation in patients with COVID-19 compared to controls (Maździarz et al., 2025). We further applied f-*ρ*PCA to transcript level expression, finding that an isoform of the gene *OAS1* known to reduce COVID-19 disease severity separates target samples better than the more common and highly expressed isoform (Zhou et al., 2021) (Supp. Fig. S5). These examples demonstrate that f-*ρ*PCA can be used to contrast two time courses of gene expression immune response in a single analysis, a complementary approach to performing post hoc comparisons of two separate analyses.

## Discussion

We have presented two extensions of the *ρ*PCA method that expand its applicability to spatial and functional data. The first extension, k-*ρ*PCA, uses kernel weighting to incorporate spatially-structured variation, and the second extension, f-*ρ*PCA, utilizes basis functions to identify contrastive modes of variation in functional data. Together, these methods show that the Rayleigh quotient formalism unifies linear contrastive analysis within a principled and tractable framework. Importantly, these extensions inherit the theoretical and practical advantages of *ρ*PCA (Carilli et al., 2025) and can be further developed, for example, by combining f-*ρ*PCA and k-*ρ*PCA.

In practice, k-*ρ*PCA is well-suited for settings where observations in a target dataset are spatially registered. A key motivation for using a kernel is that spatial patterning in standard PCA is incidental and reflects structure that informs total variance rather than spatial variance. Instead, the kernel-based method up-weights local covariance and produces components that reflect spatial structure by design. Combined with the contrastive term, k-*ρ*PCA is ideal for spatially-resolved transcriptomics data analysis where the goal is to identify tissue-specific expression patterns while controlling for normal cell-type variation. Notably, the background dataset need not be a matched sample to the target (Supp. Fig. S3), meaning that k-*ρ*PCA can leverage publicly available, non-spatial scRNA-seq data, making the method a flexible tool for integrating information from single-cell and spatially-resolved transcriptomics datasets. The approach is not limited to the modalities considered here and can, in principle, be adapted to a growing number of multimodal technologies that can characterize complex background variance distributions. Moreover, since there is established and effective machinery for computing generalized eigenvalues, k-*ρ*PCA can efficiently operate on large-scale spatial datasets, such as those produced by Visium HD.

By comparison, f-*ρ*PCA identifies modes of variation that distinguish two groups of functional observations. Crucially, f-*ρ*PCA operates in the space of basis function coefficients rather than on discrete measurements, producing eigenfunctions that are directly interpretable as temporal modes of variation. The result is that f-*ρ*PCA is well-suited for experimental designs involving paired and sequential conditions, such as dose comparisons or longitudinal case-control studies. In cases where the goal is to identify functional differences in the response between groups, f-*ρ*PCA unifies the comparative analysis rather than requiring pairwise post-hoc tests. Moreover, f-*ρ*PCA can, in principle, be extended to joint analysis of several features, incorporated as multivariate functional observations rather than treated independently. However, such an extension requires a framework for performing tensor SVD that retains the spectral properties of its lower dimensional counterparts. This remains an area of active research (Zhang and Xia, 2020).

Prior to this work, kernel, spatial, and functional PCA have existed largely as separate methodologies with distinct motivations (Shang and Zhou, 2022; Yao et al., 2005; Zhang and Li, 2025; Schölkopf et al., 1997). We note, however, that these methods share an underlying principle in that the co-variance operator can be weighted to reflect latent spatial or temporal structure. Previous work by Yao et al. (2005) implicitly makes this connection by using the estimated latent covariance function as a kernel to borrow strength across irregularly sampled time points in functional data, with the latent covariance function performing an analogous role to the kernel function in k-*ρ*PCA. The Rayleigh quotient formulation of *ρ*PCA extends this approach by providing a mechanism for performing contrastive analysis and by unifying these problems analytically, showing that they can be interpreted as the same generalized eigenvalue problem. Finally, the examples presented in this work demonstrate that both methods can recover biologically relevant signals from complex genomic datasets. We anticipate that our practical, flexible approach will find broad utility across biological domains.

## Methods

A detailed description of the *ρ*PCA method is provided in Carilli et al. (2025). Briefly, the *ρ*PCA objective finds directions that maximize variance in a target dataset relative to a background by maximizing a Rayleigh quotient. Imposing a unit-variance constraint in the background results in a constrained optimization problem that can be formulated as a generalized eigenvalue problem. The solutions to this problem are the generalized eigenvectors that define directions that maximize the target variance relative to the background variance, while their corresponding eigenvalues directly measure the target-to-background variance ratio.

### Solving k-*ρ*PCA

The k-*ρ*PCA method replaces the sample covariance matrices in the *ρ*PCA objective with their kernel-weighted counterparts with the goal of defining contrastive directions that identify latent structure encoded by the kernel. Though we prioritize cases where a spatially-indexed transcriptomic dataset is contrasted to a non-spatial scRNA-seq background, the approach inherently allows for comparison between heterogeneous combinations of indexed and non-indexed data. Further rationale is provided in the Supplementary Methods.

We consider a spatially-resolved transcriptomics experiment with measurements of *p* features across *n* spatial locations, which we denote **s**_*i*_ for *i* = 1,…, *n*. Typically, the locations are defined over a two-dimensional spatial domain, such that **s**_*i*_ = (*s*_*i*1_, *s*_*i*2_) ∈ ℝ^2^. In principle, however, the locations can be defined over an *m*-dimensional domain. The *n* × *p* matrix *X* represents the normalized and zero-centered gene expression matrix for the study.

The kernel matrix *K* is calculated from pairs of spatial coordinates, **s**_*i*_, **s**_*j*_, where each entry *K*_*ij*_ = *K*(**s**_*i*_, **s**_*j*_) represents a spatial weight between locations. A common choice of kernel function includes the Gaussian kernel:

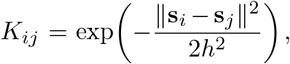

which depends on the hyperparameter *h >* 0, although alternative kernel functions have been used for other supervised learning applications (Alvarez et al., 2012). In practice, heuristic approaches are used to determine an appropriate value for the parameter *h* (Sheather and Jones, 1991). For simplicity, the default for the parameter *h* is the square root of the median pairwise distance. We therefore define the kernel-weighted sample covariance matrix, 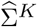, as

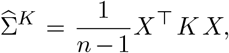

and the corresponding k-*ρ*PCA objective as

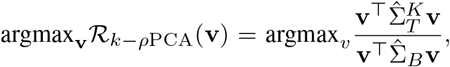

where 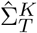 and 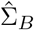 denote the empirical kernel-weighted target covariance and background covariance, respectively, and 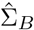 is assumed to be positive definite. The solutions **v** represent the spatial contrastive components. These vectors identify directions in the gene expression space that exhibit high spatial variance in the target tissue while remaining invariant to the non-spatial noise present in the background samples.

### Solving f-*ρ*PCA

The f-*ρ*PCA method solves the Rayleigh Quotient generalized eigenproblem for functional data. We consider *n* independent observations *X*_*i*_(*t*), …, *X*_*n*_(*t*) of some square-integrable stochastic process *X*(*t*), which we are interested in as a target process, and *m* independent observations *Y*_*i*_(*t*), …, *Y*_*m*_(*t*) of a different square-integrable stochastic process *Y* (*t*), which we treat as a background process.

Let *µ*_*Z*_(*t*) = *E*[*Z*(*t*)] and *c*_*Z*_(*s, t*) = Cov(*Z*(*s*), *Z*(*t*)) = *E*[(*Z*(*s*) −*µ*_*Z*_(*s*))(*Z*(*t*) −*µ*_*Z*_(*t*))] be the mean and covariance functions of the processes *Z*(*t*), where *Z* ∈ {*X, Y*} indicates target or background, respectively. The covariance functions are kernels for target and background covariance operators: 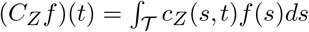. We seek to find functions *ϕ*(*t*) that maximize the Rayleigh quotient:

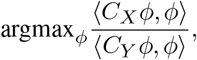

requiring that *C*_*Y*_ is positive definite on its domain, or ⟨*C*_*Y*_ *f, f*⟩ *>* 0 for non-zero functions *f* (e.g., covariance operators for non-degenerate Gaussian processes (Rasmussen and Williams, 2005)).

### Discrete domain

In practice, when functional observations have been measured at *p* discrete points, or *X* ∈ ℝ^*n×p*^ and *Y* ∈ ℝ^*m×p*^, target and background sample covariance matrices 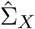 and 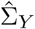 can be calculated and the discrete *ρ*PCA objective solved (Carilli et al., 2025):

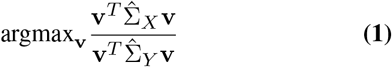

for discrete eigenvectors *v* ∈ ℝ^*p*^. These eigenvectors can be interpolated between discrete features for continuous support (Ramsay and Silverman, 2005).

### Basis representation

Alternatively, and commonly when data is irregularly or sparsely sampled, the Rayleigh quotient can be solved using a basis function representation for samples, as in functional PCA (Ramsay and Silverman, 2005). Full details are included in the Supplementary Methods ‘Basis Expansion for Functional *ρ*PCA.’ Briefly, *B*(*t*) = (*b*_1_(*t*), …, *b*_*D*_(*t*))^*T*^, a set of linearly independent basis functions, are fit to each sample:

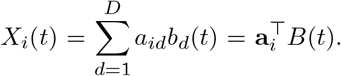

All *n* observations can be written in matrix form, *X*(*t*) = *A*_*X*_ *B*(*t*), where the matrix *A*_*X*_ ∈ ℝ^*n×D*^ contains *n* rows, one per observation, of coefficients for the *D* basis functions. The background can be similarly fit and represented as *Y* (*t*) = *A*_*Y*_ *B*(*t*). The Rayleigh quotient is then optimized to find eigenvectors **w** in coefficient space, using the bases’ Gram matrix *G* ∈ ℝ^*D×D*^ with entries *G*_*kl*_ = ∫_𝒯_ *b*_*k*_(*t*) *b*_*l*_(*t*) *dt* to account for possible non-orthogonality of functions:

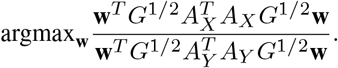

The resulting coefficient vectors **w** are then converted back to the original data space to find eigenfunctions *v*(*t*) = **w**^*T*^ *G*^1*/*2^*B*(*t*).

### Visium and Visium HD Data Processing

We obtained raw Visium V2 and Visium HD count matrices and metadata from the 10X genomics data portal (10x Genomics, 2024b). The Visium data included gene counts for 18,085 genes in 4,269 spots. For the Visium HD example, we used the count matrix associated with the 8 *µ*m segmentation which included counts for 18,085 genes in 545,913 bins.

For the Visium data, we filtered spots with fewer than 10 counts and genes expressed in fewer than 50 spots. Spots with greater than 20% mitochondrial counts were also removed. All counts were depth normalized and logtransformed. This workflow retained all spots and removed 50 genes from the subsequent analysis. To process the Visium HD data, we followed the workflow described by the original authors in the resource provided at (10x Genomics, 2025). Briefly, genes expressed in fewer than 50 bins were excluded from further analysis. Additionally, bins with fewer than 53 counts and more than 2,979 counts were removed. After processing, 471,258 bins and 16,946 genes were available for analysis.

### Application of k-*ρ*PCA to Visium and Visium HD

For both the Visium and Visium HD datasets, we filtered to the 3,000 most highly-variable genes identified in the matched scRNA-seq data from Patient 2 as background. We obtained scRNA-seq counts from (10x Genomics, 2024a) to use an unmatched background in Supplementary Figure S3, and used the same filtering procedure to generate the components. In the case of the Visium HD data, we used a radius-based approach to define spatial neighbors and chose the radius such that an average of 800 spatial neighbors were available for each bin.

### Longitudinal Bulk RNA-seq Data Processing

We obtained the aligned bulk RNA-seq counts per sample and time point for booster and primer doses (pre-normalization, see original publication for details) from (Rinchai et al., 2022) from the NCBI GEO repository under accession ID GSE190001. This longitudinal study sequenced blood transcriptomes from 23 patients immediately before, for 9 days after, and 14 days after vaccination with a primer and booster COVID-19 mRNA vaccine. The primer matrix included 23 unique patients and 11 unique time points for a total 213 samples of 58,735 genes (not all patients were collected at each time point). The booster dataset included the same 23 unique patients and 11 unique time points for a total 226 samples of 58,735 genes.

Following the original study, we normalized primer and booster counts using `pydeseq2 0.5.4`, then subset to the 119 interferon genes of ‘module A28’ reported in (Rinchai et al., 2022). We then normalized each gene expression trajectory per patient for booster and primer time courses (separately) by dividing by the value pre-dose (day 0).

### Application of f-*ρ*PCA to Longitudinal Bulk RNA-seq

For each of the 119 genes, we fit patients’ normalized trajectories to a B-spline basis with 5 splines using `skfda.representation.basis.BSplineBasis` with `skfda` version 0.10.1 (Ramos-Carreño et al., 2024). We kept trajectories with an *R*^2^ of at least 0.5. For genes with at least 7 primer and at least 7 booster trajectories that passed the *R*^2^ threshold (86 genes), we performed f-*ρ*PCA with booster trajectories as target and primer trajectories as background in the basis representation to find the top eigenfunctions per gene. For each gene, we projected booster and primer trajectories onto the top eigenfunction to obtain f-GE1 scores, and calculated the ratio of variance of booster f-GE1 scores to variance of primer f-GE1 scores, allowing us to order genes by their target-to-background projected variance ratio.

## Code and data availability

The *ρ*PCA suite of tools is available as a pip installable package at https://github.com/pachterlab/rhopca/. Code and data for reproducing manuscript results can be found at https://github.com/pachterlab/CJP_2025/.

All data used in this study are available from the original authors. The Visium and Visium HD colorectal cancer data were obtained from 10X Genomics at https://www.10xgenomics.com/platforms/visium/product-family/dataset-human-crc. The colorectal scRNA-seq data from 10X Genomics was obtained from https://www.10xgenomics.com/datasets/320k_scFFPE_16-plex_GEM-X_FLEX. The aligned bulk RNA-seq counts from Rinchai et al. (2022) are available from the NCBI GEO repository under accession ID GSE190001.

## Acknowledgments

Thanks to members of the Pachter lab who provided valuable input in discussions related to the methods during the journal club that led to the work on this paper. MC was funded, in part, by the NSF GRFP under grant no. 2139433. KJ and LP were funded, in part, by NIH 1R01DK143671-02.

## Author contributions

This work is a companion to (Carilli et al., 2025). KJ, MC, and LP developed the methods, produced the results and drafted the manuscript.

## Notes

### Competing Interest Statement

The authors have declared no competing interest.

