## Supplementary_Figures for "The Rayleigh Quotient and Contrastive Principal Component Analysis II"

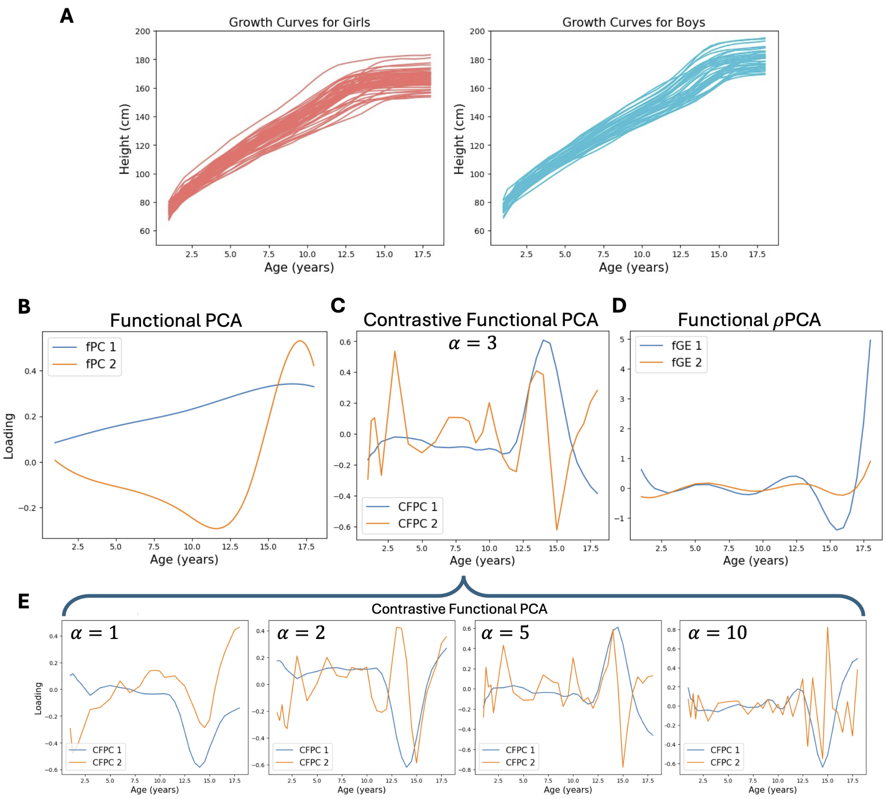


Figure S1: Comparison of functional PCA [Wang et al., 2016], contrastive functional PCA [Zhang and Li, 2025], and functional $\rho$PCA on the Berkeley height dataset [Tuddenham and Snyder, 1954]. **A.** Measured growth curves of the height (in cm) of n = 54 girls (orange, left) and n = 39 boys (blue, right) from [Tuddenham and Snyder, 1954]. **B.** The first two functional principal components fit using 7 B-spline bases on all individuals (boys and girls). **C.** The first two contrastive functional principal components [Zhang and Li, 2025] with contrastive parameter $\alpha = 3$ using the boys' heights as foreground and the girls' heights as background. **D.** First two generalized eigenfunctions found using functional $\rho$PCA (f-$\rho$PCA) fit using 7 B-spline bases with the boys' heights as foreground and the girls' heights as background. **E.** Contrastive functional PCA with four different contrastive parameters ($\alpha$ = 1, 2, 5, and 10) result in different first and second functional principal components. See Supplementary Methods for fit details.

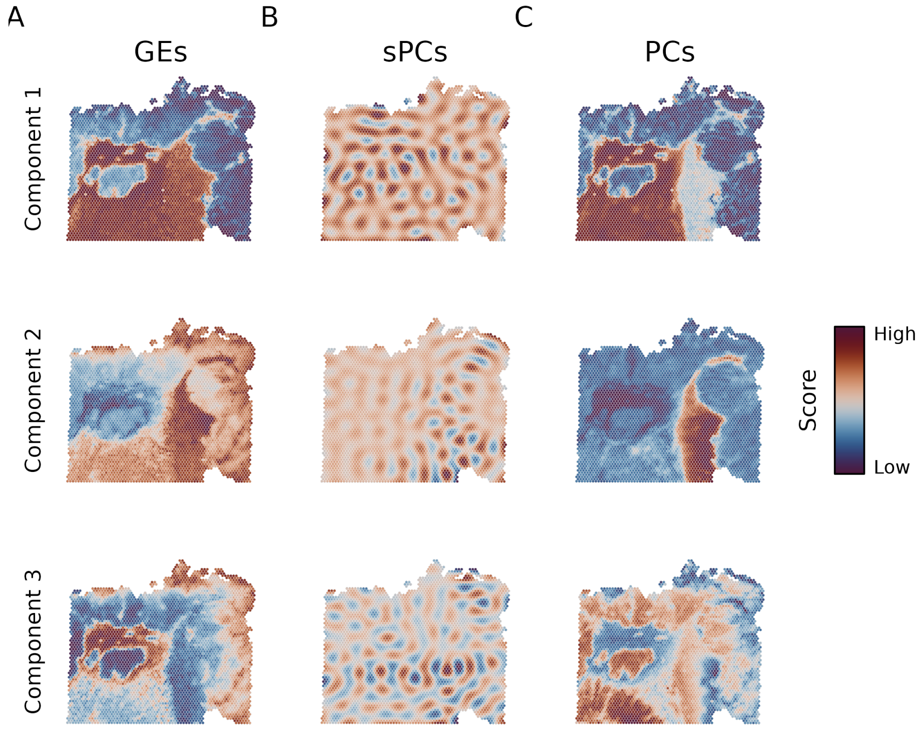


Figure S2: Comparison of k-$\rho$PCA, spatial PCA [Shang and Zhou, 2022], and PCA on Visium CRC data. Color indicates the projection score of each spot onto the component indicated on the left. Both **A.** k-$\rho$PCA and **C.** PCA produce components with spatially coherent structure, while **B.** spatial PCA yields components that do not correspond to biologically-interpretable boundaries. We ran spatial PCA with the "fast" option enabled and kept other parameters set to their defaults. Spatial PCA exited with an unexplained error when we set the bandwidth to the same value used to generate the results in panel **A.**

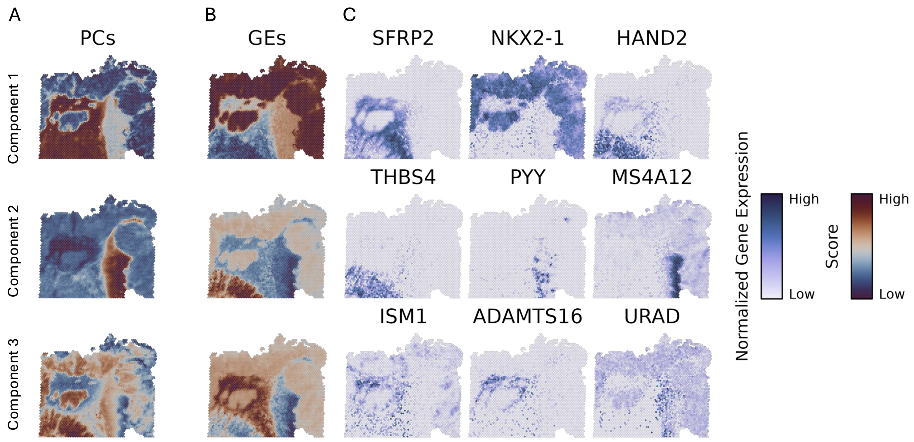


Figure S3: Application of k-$\rho$PCA to Visium V2 data using unmatched scRNA-seq as background. **A.** The PCs as in Figure S2. **B.** The GEs and their top genes (**C.**) computed from the Visium V2 target and unmatched single-cell FFPE dataset downloaded from 10X Genomics.

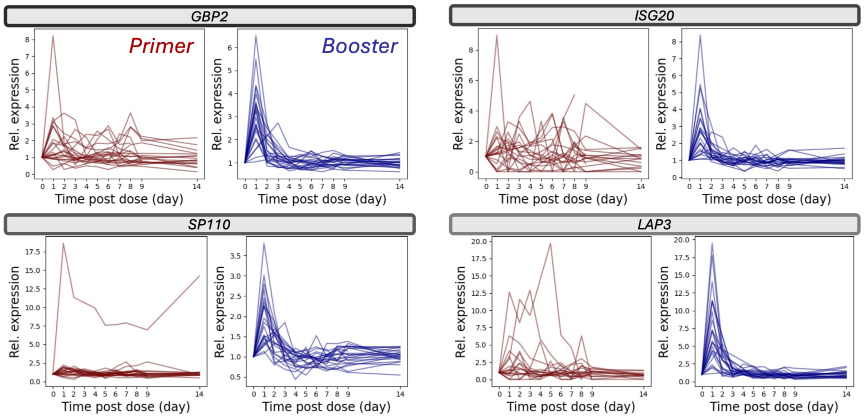


Figure S4: Gene expression measured using bulk RNA-seq over time following a first ("primer") and second ("booster") COVID-19 mRNA vaccine dose (from [Rinchai et al., 2022]). Each curve is a separate patient, and each is normalized to the initial time point (see Main Methods for data processing). The genes (*GBP2*, with n = 10 primer samples and n = 7 booster samples passing goodness of fit criteria; *ISG20*, with n = 8 primer and n = 8 booster samples passing goodness of fit criteria; *SP110*, with n = 11 primer and n = 12 booster samples passing goodness of fit criteria; and *LAP3*, with n = 7 primer and n = 7 booster samples passing goodness of fit criteria) are those along which the variance of booster to primer projections was greatest (see Main Fig. 2C).

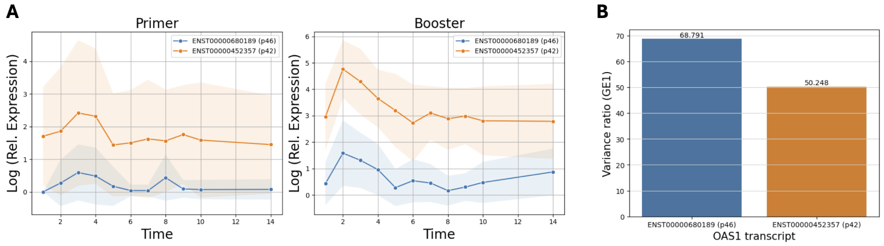


Figure S5: Bulk RNA-seq data from a longitudinal study on blood samples following COVID-19 mRNA vaccination [Rinchai et al., 2022] (processing and fit details in Supplementary Methods) demonstrate that f-$\rho$PCA can be used to investigate isoform level differences in expression profiles. **A.** The log of normalized transcript expression for the two most common *OAS1* transcripts over 23 patients (mean line and shaded standard deviation for ENST00000680189, or isoform p46, in blue and ENST00000452357, or isoform p42, in orange) following a primer and booster dose. **B.** The variance ratio of booster to primer patients projected onto the first generalized eigenfunction is higher for p46 isoform, which is known to reduce COVID-19 severity [Zhou et al., 2021].

### References

Darawan Rinchai, Sara Deola, Gabriele Zoppoli, Basirudeen Syed Ahamed Kabeer, Sara Taleb, Igor Pavlovski, Selma Maacha, Giusy Gentilcore, Mohammed Toufiq, Lisa Mathew, Li Liu, Fazulur Rehaman Vempalli, Ghada Mubarak, Stephan Lorenz, Irene Sivieri, Gabriella Cirmena, Chiara Dentone, Paola Cuccarolo, Daniele Roberto Giacobbe, Federico Baldi, Alberto Garbarino, Benedetta Cigolini, Paolo Cremonesi, Michele Bedognetti, Alberto Ballestrero, Matteo Bassetti, Boris P. Hejblum, Tracy Augustine, Nicholas Van Panhuys, Rodolphe Thiebaut, Ricardo Branco, Tracey Chew, Maryam Shojaei, Kirsty Short, Carl G. Feng, PREDICT-19 Consortium, Susu M. Zughaier, Andrea De Maria, Benjamin Tang, Ali Ait Hssain, Davide Bedognetti, Jean-Charles Grivel, and Damien Chaussabel. High–temporal resolution profiling reveals distinct immune trajectories following the first and second doses of covid-19 mrna vaccines. *Science Advances*, 8(45), 2022. doi: 10.1126/sciadv.abp9961.

L. Shang and X. Zhou. Spatially aware dimension reduction for spatial transcriptomics. *Nature Communications*, 13(1):7203, Nov 2022. doi: 10.1038/s41467-022-34879-1.

Robert D. Tuddenham and Margaret M. Snyder. Physical growth of california boys and girls from birth to age 18. *California Publications in Child Development*, 1:183–364, 1954.

Jane-Ling Wang, Jeng-Min Chiou, and Hans-Georg Müller. Functional data analysis. *Annual Review of Statistics and Its Application*, 3:257–295, 2016. doi: 10.1146/annurev-statistics-041715-033624.

Eric Zhang and Didong Li. Contrastive functional principal component analysis. In *Proceedings of the AAAI Conference on Artificial Intelligence*, volume 39, pages 22380–22388, 2025.

Sirui Zhou, Guillaume Butler-Laporte, Tomoko Nakanishi, David R. Morrison, Jonathan Afilalo, Marc Afilalo, Laetitia Laurent, Maik Pietzner, Nicola Kerrison, Kaiqiong Zhao, Elsa Brunet-Ratnasingham, Danielle Henry, Nofar Kimchi, Zaman Afrasiabi, Nardin Rezk, Meriem Bouab, Louis Petitjean, Charlotte Guzman, Xiaoqing Xue, Chris Tselios, Branka Vulesevic, Olumide Adeleye, Tala Abdullah, Noor Almamlouk, Yiheng Chen, Michaël Chassé, Madeleine Durand, Clare Paterson, Johan Normark, Robert Frithiof, Miklós Lipcsey, Michael Hultström, Celia M. T. Greenwood, Hugo Zeberg, Claudia Langenberg, Elin Thysell, Michael Pollak, Vincent Mooser, Vincenzo Forgetta, Daniel E. Kaufmann, and J. Brent Richards. A neanderthal oas1 isoform protects individuals of european ancestry against covid-19 susceptibility and severity. *Nature Medicine*, 27:659–667, 2021. doi: 10.1038/s41591-021-01281-1.
