## Supplementary_Methods for "The Rayleigh Quotient and Contrastive Principal Component Analysis II"

### Rationale for kernel $\rho$PCA

Spatial applications of kernel $\rho$-PCA (k-$\rho$PCA) analyze a covariance matrix that encodes spatial relationships from a pairwise spatial kernel. Kernel information is stored in the matrix $\mathbf{K}$ which contains positive weights for pairs of observations. Most commonly, the entries in $\mathbf{K}$, $\mathbf{K}_{ij} = K(x_i, x_j)$, are the output of a positive semi-definite (PSD) kernel that operates on each pair of observations.

The $n \times p$ expression matrix $X$ contains gene expression information of the $p$ features across $n$ observations. We consider the features in $X$ as centered and weakly stationary spatial processes, where $X_i = \{X_i(s) : s \in \mathbb{R}^d\}$ for $i = 1, \ldots, p$. With these assumptions, the covariance between features can be estimated directly:

$$\begin{aligned}
\mathrm{Cov}\big(X_i(S), X_j(S')\big) &= \mathrm{E}\big[\mathrm{Cov}\big(X_i(S), X_j(S') \mid S, S'\big)\big] \\
&\quad + \mathrm{Cov}\big(\mathrm{E}\big[X_i(S) \mid S, S'\big], \mathrm{E}\big[X_j(S') \mid S, S'\big]\big) \\
&= \mathrm{E}\big[\mathrm{Cov}\big(X_i(S), X_j(S') \mid S = s, S' = s'\big)\big] \\
&= \mathrm{E}\big[\mathrm{Cov}\big(X_i(s), X_j(s')\big)\big] \\
&= \mathrm{E}\big[C_{X_i X_j}(s, s')\big]\,,
\end{aligned}$$

where

$$C_{X_i X_j}(s, s') \;=\; \mathrm{Cov}\big(X_i(s), X_j(s')\big)$$

is the cross covariance function, which k-$\rho$PCA models as

$$C_{XY}(s, s') \;=\; \mathrm{E}[X(s)Y(s')]K(s, s')$$

In practice, however, there is only one realization of the spatial process and the covariance is estimated by averaging products across spatial pairs. Thus, the sample kernel-weighted covariance can be written compactly as

$$\hat{\Sigma}^K \;\propto\; X^\top K X$$

For other applications, the kernel matrix can be described in terms of inner products for a given feature mapping $\phi$. More specifically,

$$K_{ij}(\mathbf{x}_i, \mathbf{x}_j) \;=\; \phi(\mathbf{x}_i)^\top \phi(\mathbf{x}_j)$$

Mercer's theorem guarantees that any valid kernel function produces a symmetric, positive semi-definite matrix. This result is precisely the Gram matrix generated by the feature mappings $\phi(\mathbf{x}_1) \ldots \phi(\mathbf{x}_n)$ where $G_{ij} = \big\langle \phi(\mathbf{x}_i), \phi(\mathbf{x}_j) \big\rangle = K_{ij}$. Thus, computing the kernel-weighted covariance matrix, $X^\top K\, X$,with centered data $X$, maps $K$ to the projective space of centered Gram matrices. This ensures that the principal components of the weighted matrix are invariant to global shifts in the kernel function. To see this, we first consider a constant matrix $C = c\mathbf{1}\mathbf{1}^\top$. Using the shifted kernel-matrix to compute the weighted covariance we have:

$$
\begin{aligned}
X^\top (C + G)\, X &= X^\top \big(c\mathbf{1}\mathbf{1}^\top + G\big)\, X \\
&= X^\top \big(c\mathbf{1}\mathbf{1}^\top\big)\, X + X^\top G\, X \\
&= c\big(X^\top \mathbf{1}\big)\big(\mathbf{1}^\top X\big) + X^\top G\, X \\
&= c\big(\mathbf{0}\big)\big(\mathbf{0}^\top\big) + X^\top G\, X \\
&= X^\top G\, X
\end{aligned}
$$

Consequently, any kernel and its shifted counterpart represent identical configurations of the data in the new feature space and share spectral properties, including eigenvectors. It is also straightforward to show that scaling the kernel matrix by a constant scalar $c$ results in the same solution with eigenvalues scaled by $c$.

#### Comparison to spatial PCA (Shang and Zhou, 2022)

For the comparison to spatial PCA (Shang and Zhou, 2022) in Supplementary Figure S2, we filtered the Visium data to the same set of genes that we used in the k-$\rho$PCA analysis, and used the "fast" option to generate the spatial components. We were unable to run the method to completion without the "fast" option. Other arguments were kept at the recommended defaults when running spatial PCA, including the bandwidth for the Gaussian kernel. When we used the same bandwidth that we used to generate the results in k-$\rho$PCA, we found that the program exited with error unless we changed other default options. Thus, we chose to use the default bandwidth settings and the default argument choices.

#### Basis expansion for functional $\rho$PCA

Functional PCA (fPCA) is an extension of principal component analysis for functional data that reduces the dimension of infinite-dimensional observations of functions, with each observation considered as an independent realization of a stochastic process (Wang et al., 2016). As data are inherently infinite dimensional, reducing data to discover key "modes of variation" is an important step in analysis (Rice and Silverman, 1991). In practice, experimental data are collected on discrete fixed or random time grids, and various methods have been proposed to smooth curves and estimate mean and covariance functions over time (Yao et al., 2005). A popular implementation of fPCA first transforms data to a basis representation and uses the resulting coefficients to obtain coefficients for eigenfunctions in the basis space (Ramos-Carreño et al., 2024) that, among all basis expansions of the same number of basis functions, maximizes the variance in the data. This approach can be adapted to find generalized eigenfunctions that describe modes of variation present in a sample of target curves and not in background curves.

First, let $n$ independent mean-centered trajectories $X_i(t), i = 1, ..., n$ be represented on a basis of $D$ functions (by fitting discrete observation points per trajectory to a B-spline, Fourier or other basis (Ramsay and Silverman, 2005; Gertheiss et al., 2024)):

$$X_i(t) = \sum_{d=1}^{D} a_{id} b_d(t) \; = \; \mathbf{a}_i^\top B(t),$$
$$B(t) = (b_1(t), ..., b_D(t))^T,$$

where $B(t)$ are linearly independent basis functions. All observations can be written in matrix form, $X(t) = AB(t)$, where the matrix $A \in \mathbb{R}^{n \times D}$ contains rows per observation of coefficients for the $D$ basis functions. The Gram matrix $G \in \mathbb{R}^{D \times D}$ for the basis $B(t)$ is a matrix of inner products between basis functions over their domain $[t_0, t_f]$:

$$G_{ij} \; = \int_{t_0}^{t_f} b_i(t) b_j(t) \, dt,$$

where the time argument in $B(t)$ has been omitted for shorthand. As $G$ is positive semi-definite, it admits a symmetric square root. Let $L := G^{1/2}$ so that $G = LL^\top$.

Now, given a mean-centered target dataset $X(t) = A_X B(t)$ and a mean-centered background set of curves represented in the same basis, $Y(t) = A_Y B(t)$, we would like to find coefficient vectors $\mathbf{u}_k, k = 1, ..., K$ in the basis $B(t)$ that maximize variance in $X$ while minimizing it in $Y$. To account for non-orthogonality of basis functions, we first transform the coefficient space using the square root of the associated Gram matrix ($G^{1/2} = L$).

Let $\Sigma_X := X^T X = B^\top A_X^\top A_X B$ and $\Sigma_Y := Y^\top Y = B^\top A_Y^\top A_Y B$ be the data covariance matrices, and set $F_X := A_X G L^{-\top}, F_Y := A_Y G L^{-\top}$. The generalized eigenproblem

$$F_X^\top F_X \mathbf{w} \;=\; \lambda F_Y^\top F_Y \mathbf{w}$$

can be written as

$$L^{-1}\; G^\top\; A_X^\top\; A_X\; G\; L^{-\top}\; \mathbf{w} \;=\; \lambda\; L^{-1}\; G^\top\; A_Y^\top\; A_Y\; G\; L^{-\top}\; \mathbf{w}.$$

Using the fact that $G = G^\top = BB^\top$, we obtain

$$L^{-1}\; B\; B^\top\; A_X^\top\; A_X\; B\; B^\top\; L^{-\top}\; \mathbf{w} \;=\; \lambda\; L^{-1}\; B\; B^\top\; A_Y^\top\; A_Y\; B\; B^\top\; L^{-\top}\; \mathbf{w},$$

which, when written in terms of the data covariance matrices, is

$$L^{-1}\; B\; \Sigma_X\; B^\top\; L^{-\top}\; \mathbf{w} \;=\; \lambda\; L^{-1}\; B\; \Sigma_Y\; B^\top\; L^{-\top}\; \mathbf{w}.$$

Defining $\mathbf{u} := L^{-\top}\mathbf{w}$ and left-multiplying by $B^{-1}L$ yields

$$\Sigma_X B^\top \mathbf{u} \;=\; \lambda \Sigma_Y B^\top \mathbf{u}.$$

Thus, the generalized eigenfunctions are given by

$$\phi(t) \;=\; \mathbf{u}^\top B(t),$$

where the eigenvectors $\mathbf{u}$ are the coefficient vectors of the generalized eigenfunctions expressed in the original basis $B(t)$. In practice, we solve the generalized eigenproblem $F_X^\top F_X \mathbf{w} \;=\; \lambda F_Y^\top F_Y \mathbf{w}$ to obtain $\mathbf{w}$, and then recover the eigenfunctions as basis function coefficients via the transformation $\mathbf{u} = L^{-\top}\mathbf{w}$.

##### f-$\rho$PCA on Berkeley Height Dataset

To compare functional PCA, contrastive functional PCA, and functional $\rho$PCA (see Fig. S2) we used longitudinal height measurements from the Berkeley height study (Tuddenham and Snyder, 1954). Data from the Berkeley height study was obtained using `scikit-fda` method `datasets.fetch_growth` (Ramos-Carreño et al., 2024). This includes 31 height measurements (in cm) of 39 boys and 54 girls from the ages of 1 to 18 years. The previous and all following functions from `scikit-fda` (`skfda`) were version 0.10.1. Using a B-Spline basis (`skfda.representation.basis.BSplineBasis`) with 7 basis functions, we fit a functional PCA (`skfda.preprocessing.dim_reduction.FPCA`) with 2 components. We then fit contrastive functional PCA (CFPCA) on the data using their main function `CFPCA` with several different contrastive parameters: $\alpha = 1, 2, 5,$ and 10 (Zhang and Li, 2025). We then fit functional $\rho$-PCA using a B-Spline representation with 7 basis functions to find the first two contrastive eigenfunctions. Supplementary Figure S2 shows eigenfunction results of the fits.

##### f-$\rho$PCA on *OAS1* isoforms

To demonstrate the effectiveness of f-$\rho$PCA to discover transcript level profile differences, we set out to investigate isoform usage of *OAS1* following a first and second dose of the COVID-19 mRNA vaccine (Rinchai et al., 2022). The two most common isoforms of *OAS1* are p46 and p42: the p46 isoform decreases COVID-19 susceptibility and severity and its higher expression is associated with the Neanderthal genetic variant rs10774671 (Zhou et al., 2021).

We obtained raw FASTQ files from NCBI GEO repository under accession ID GSE190001 (Rinchai et al., 2022). This contained sequenced blood transcriptomes from 23 patients in a 14 day time period before and after COVID-19 primer and booster doses (a total of 213 primer samples and 226 booster samples). We aligned the raw data using `kallisto 0.50.1` and quantified with `kb-python 0.28.2` to find transcript level expression (Bray et al., 2016; Melsted et al., 2021). We then normalized raw counts per sample by the TMM (Trimmed Mean of M-values (Robinson et al., 2010)) and then further normalized each patient's booster and primer gene trajectories to the expression of that gene at day 0 for the booster and primer time courses, respectively. We then performed f-$\rho$PCA on the most highly expressed transcript for the p46 isoform (ENST00000680189) and the p42 (ENST00000452357) isoforms using booster samples as the target and primer samples as the background. We first fit all samples to a B-Spline basis with 5 basis functions (`skfda.representation.basis.BSplineBasis`) (Pedregosa et al., 2011) and performed f-$\rho$PCA on the coefficients as described in the Main Methods and Supplementary Note 'Basis expansion for functional $\rho$PCA.' We projected samples onto the first generalized eigenfunction for both transcripts and calculated the ratio of the variance of target to the variance of background samples, shown in Supp. Fig. S5.

Darawan Rinchai, Sara Deola, Gabriele Zoppoli, Basirudeen Syed Ahamed Kabeer, Sara Taleb, Igor Pavlovski, Selma Maacha, Giusy Gentilcore, Mohammed Toufiq, Lisa Mathew, Li Liu, Fazulur Rehaman Vempalli, Ghada Mubarak, Stephan Lorenz, Irene Sivieri, Gabriella Cirmena, Chiara Dentone, Paola Cuccarolo, Daniele Roberto Giacobbe, Federico Baldi, Alberto Garbarino, Benedetta Cigolini, Paolo Cremonesi, Michele Bedognetti, Alberto Ballestrero, Matteo Bassetti, Boris P. Hejblum, Tracy Augustine, Nicholas Van Panhuys, Rodolphe Thiebaut, Ricardo Branco, Tracey Chew, Maryam Shojaei, Kirsty Short, Carl G. Feng, PREDICT-19 Consortium, Susu M. Zughaier, Andrea De Maria, Benjamin Tang, Ali Ait Hssain, Davide Bedognetti, Jean-Charles Grivel, and Damien Chaussabel. High–temporal resolution profiling reveals distinct immune trajectories following the first and second doses of covid-19 mrna vaccines. *Science Advances*, 8(45), 2022. doi: 10.1126/sciadv.abp9961.

Mark D. Robinson, Davis J. McCarthy, and Gordon K. Smyth. edger: a bioconductor package for differential expression analysis of digital gene expression data. *Bioinformatics*, 26(1):139–140, 2010. doi: 10.1093/bioinformatics/btp616.

L. Shang and X. Zhou. Spatially aware dimension reduction for spatial transcriptomics. *Nature Communications*, 13(1):7203, Nov 2022. doi: 10.1038/s41467-022-34879-1.

Robert D. Tuddenham and Margaret M. Snyder. Physical growth of california boys and girls from birth to age 18. *California Publications in Child Development*, 1:183–364, 1954.

Jane-Ling Wang, Jeng-Min Chiou, and Hans-Georg Müller. Functional data analysis. *Annual Review of Statistics and Its Application*, 3:257–295, 2016. doi: 10.1146/annurev-statistics-041715-033624.

Fang Yao, Hans-Georg Müller, and Jane-Ling Wang. Functional data analysis for sparse longitudinal data. *Journal of the American statistical association*, 100(470):577–590, 2005.

Eric Zhang and Didong Li. Contrastive functional principal component analysis. In *Proceedings of the AAAI Conference on Artificial Intelligence*, volume 39, pages 22380–22388, 2025.

Sirui Zhou, Guillaume Butler-Laporte, Tomoko Nakanishi, David R. Morrison, Jonathan Afilalo, Marc Afilalo, Laetitia Laurent, Maik Pietzner, Nicola Kerrison, Kaiqiong Zhao, Elsa Brunet-Ratnasingham, Danielle Henry, Nofar Kimchi, Zaman Afrasiabi, Nardin Rezk, Meriem Bouab, Louis Petitjean, Charlotte Guzman, Xiaoqing Xue, Chris Tselios, Branka Vulesevic, Olumide Adeleye, Tala Abdullah, Noor Almamlouk, Yiheng Chen, Michaël Chassé, Madeleine Durand, Clare Paterson, Johan Normark, Robert Frithiof, Miklós Lipcsey, Michael Hultström, Celia M. T. Greenwood, Hugo Zeberg, Claudia Langenberg, Elin Thysell, Michael Pollak, Vincent Mooser, Vincenzo Forgetta, Daniel E. Kaufmann, and J. Brent Richards. A neanderthal oas1 isoform protects individuals of european ancestry against covid-19 susceptibility and severity. *Nature Medicine*, 27:659–667, 2021. doi: 10.1038/s41591-021-01281-1.
